# A Multiscale Computational Framework for the ^28^Mg Radio-Cofactor Hypothesis: Conditional Emergence of Coordinated Disruption under the Gate Condition

**DOI:** 10.64898/2026.08.01.742251

**Authors:** Tran Van Luyen

## Abstract

Cancer therapy continues to confront molecular redundancy, metabolic plasticity and multiscale adaptability that limit durable responses. Most existing modalities act on downstream products, signaling pathways or extracellular recognition structures, while the deeper intracellular regulatory architecture that sustains malignant proliferation remains comparatively underexplored. Enzymatic cofactors occupy a uniquely fundamental position within this architecture: they enable catalytic activity itself. The Radio-Cofactor Hypothesis proposes that an essential biological cofactor can serve as an endogenous carrier of radionuclide activity. Using magnesium-28 (^28^Mg) as prototype, the hypothesis posits that a radioactive isotope chemically indistinguishable from physiological Mg^2+^ can occupy magnesium-dependent catalytic sites; subsequent nuclear transformation then generates simultaneous alteration of cofactor identity and highly localized energy deposition within the active site.

The present study does not experimentally demonstrate catalytic-site occupancy. Instead it treats non-zero fractional occupancy (θ_28_ > 0) as an explicit input premise—the Gate Condition—and constructs a hierarchical computational discovery platform that integrates nuclear-decay physics, magnesium enzymology, intracellular transport, mitochondrial and nuclear responses, radiobiology, pharmacokinetics and tumor-growth dynamics. Information propagates across six organizational levels according to defined bottom-up and top-down rules. Under the gate-condition assumption the framework generates a sequence of emergent behaviors: the Atomic Switch / Decay-Induced Octahedral Collapse at the molecular scale, progressive Enzyme Disruption Index (EDI) across functional enzyme classes, a coordinated Quadruple-Kill cascade linking catalytic, radiolytic, mitochondrial and transcriptional injury, and a system-level Quintax Functional Model. Tissue-scale trajectories are described by an intrinsic Gompertz formulation, while whole-body dosimetry is evaluated against QUANTEC constraints.

All higher-scale predictions remain strictly conditional upon satisfaction of the gate condition and upon the phenomenological transport and uptake parameters assigned to the model. The framework is therefore hypothesis-generating rather than predictive of clinical efficacy. Its principal contribution is to convert the radio-cofactor concept into a quantitatively linked, experimentally addressable cascade and to provide a clear roadmap of decision points—beginning with verification of differential magnesium transport and catalytic-site occupancy —for systematic empirical interrogation.

## I. Introduction

Cancer therapy has advanced from non-specific cytotoxicity toward increasingly precise molecular, cellular, and immune-based interventions. Receptor-targeted agents, monoclonal antibodies, immune-checkpoint modulators, gene-directed strategies, and targeted radionuclide therapies have improved selectivity and clinical outcomes. Nevertheless, durable responses remain limited. Molecular redundancy, metabolic plasticity, and multiscale adaptability continue to drive therapeutic resistance. Most current approaches act on downstream molecular products, signaling pathways, or extracellular recognition structures, while the deeper intracellular regulatory architecture that sustains malignant proliferation remains comparatively underexplored.

Within this architecture, enzymatic cofactors occupy a uniquely fundamental position. Unlike receptors or signaling proteins that transmit information, cofactors enable catalytic activity itself. Magnesium (Mg^2+^) is one of the most abundant and versatile biological metal cofactors [1,2], participating in ATP utilization, nucleic-acid synthesis, kinase regulation, mitochondrial bioenergetics, ribosomal function, and hundreds of enzymatic reactions required for cellular survival and proliferation. If genes encode biological information and proteins execute biological functions, enzymatic cofactors determine whether those functions can occur. Despite this central role, cofactors have rarely been investigated as direct therapeutic control nodes in oncology.

The Radio-Cofactor Hypothesis [3] proposes a distinct conceptual framework in which an essential biological cofactor itself serves as an endogenous carrier of radionuclide activity. Using Magnesium-28 (^28^Mg) as a prototype, the hypothesis posits that a radioactive isotope chemically indistinguishable from physiological Mg^2+^ can participate in endogenous magnesium-dependent pathways. Once incorporated into the active sites of magnesium-dependent enzymes, subsequent nuclear transformation (^28^Mg → ^28^Al → ^28^Si) generates an “Atomic Switch”: simultaneous alteration of cofactor identity and deposition of highly localized radiation energy within the catalytic environment. This concept links enzymology, nuclear physics, and intracellular radiobiology in a manner distinct from conventional carrier-based radionuclide delivery.

A critical prerequisite underlies the entire Radio-Cofactor Hypothesis. Because ^28^Mg is chemically identical to physiological Mg^2+^ prior to decay, its ability to occupy magnesium-dependent catalytic sites is governed by the same thermodynamic binding affinities. Occupancy of any given active site is described by a competitive binding isotherm of hyperbolic form, mathematically analogous to the Michaelis–Menten equation [4,5]:

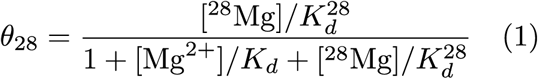

where *θ*_28_ is the fractional occupancy by ^28^Mg.

Magnesium is an essential cofactor for more than 600 enzymes and an activator for approximately 200 additional enzymes, participating in ATP utilization, nucleic-acid synthesis and repair, kinase regulation, mitochondrial bioenergetics and numerous other processes required for cellular proliferation [6,7]. If *θ*_28_ = 0 (i.e., ^28^Mg fails to compete successfully for catalytic sites), the radio-cofactor concept terminates at the molecular level and no downstream therapeutic mechanism can arise. Conversely, if non-zero occupancy is achievable, nuclear decay within the active site becomes a plausible source of multi-level biological disruption.

The present work does not experimentally demonstrate this binding step. Instead, it treats successful catalytic-site occupancy (*θ*_28_ > 0) as an explicit input premise—the gate condition —and constructs a multiscale computational framework to explore the mechanistic consequences that would follow if the gate condition is satisfied.

Because atomic-scale nuclear events must propagate across molecular, organellar, cellular, tissue, and organismal levels to produce emergent biological effects, a reductionist analysis is insufficient. We therefore developed a hierarchical computational discovery platform that integrates nuclear-decay physics, magnesium enzymology, intracellular transport, mitochondrial and nuclear responses, radiobiology, pharmacokinetics, and tumor-growth dynamics. The framework is not intended as a predictive model of clinical efficacy. Its purpose is to examine whether the integration of independently established physical and biological principles can generate previously unrecognized higher-order mechanisms and experimentally testable predictions.

Through this approach we identify candidate emergent behaviors, including the Quadruple-Kill cascade—a coordinated multi-level disruption of enzymatic catalysis, mitochondrial integrity, genomic stability, and proliferative signaling—and the Quintax Functional Model describing system-level control of radio-cofactor activity. These constructs are computationally derived mechanistic hypotheses, not experimentally confirmed phenomena. Their validity ultimately depends on systematic empirical interrogation. By clarifying the logical structure of the radio-cofactor concept and providing a quantitative roadmap for its evaluation, the framework aims to reduce mechanistic uncertainty and accelerate the transition from theoretical proposal to rigorous experimental testing.

## II. Methods

### II.1. Framework Architecture and Scope

We developed a multiscale mechanistic computational framework to investigate the biological consequences of the ^28^Mg radio-cofactor concept under the explicit premise that ^28^Mg can achieve non-zero occupancy of magnesium-dependent enzyme active sites (the gate condition). The framework connects six organizational levels—atomic/nuclear, molecular, organelle, cellular, tissue, and whole-body—through defined information-transfer rules. Each level receives quantitative or semi-quantitative inputs from the level below and supplies constrained outputs to the level above.

The architecture is conditional: all higher-scale results exist only if the gate condition is met. The framework does not compute or validate the absolute probability of ^28^Mg binding; it maps the downstream consequences once occupancy is assumed.

Detailed numerical parameters, validation logs, and executable protocols are provided in Supplementary Information.

### II.2. Cross-scale Computational Orchestration

Information propagates according to two complementary principles:

- **Bottom-up:** lower-scale physical or biochemical quantities are converted into effective parameters that drive the next higher scale.
- **Top-down:** higher-scale constraints (resource limitation, tissue carrying capacity, systemic clearance) modulate the effective rates of lower-scale processes.

The degree of quantification decreases with biological complexity:

| Interface | Nature of coupling | Primary transferred quantities |
| --- | --- | --- |
| Atomic/Nuclear → Molecular | Quantitative | Local energy density, recoil energy, Auger spectra |
| Molecular → Organelle | Semi-quantitative | Enzyme Disruption Index (EDI), catalytic-capacity modifiers |
| Organelle → Cellular | Phenomenological + scaling | Relative replication rate, metabolic fitness, survival probability |
| Cellular → Tissue | Semi-quantitative (derived) | Effective growth rate $\lambda_{eff}$, accessibility-modified $K_{eff}$ |
| Tissue → Whole-body | Quantitative (dosimetry) + phenomenological (uptake) | Time-integrated activity, absorbed dose, tumor-burden trajectory |

This graded scheme preserves mechanistic interpretability while acknowledging current limits of biological parameterization.

### II.3. Atomic/Nuclear Layer

#### Nuclear data for the sequential decay

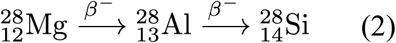

were taken from NNDC/ENSDF 2024 [8]. Decay kinetics were solved analytically with the Bateman equations (atom-conservation error < 10*^−^*^13^). Emitted *β* particles, Auger electrons, and nuclear-recoil ions (up to ∼ 414 eV for ^28^Si) were treated as nanoscale energy-deposition events. Monte-Carlo radiation-transport calculations supplied spatially resolved energy-density maps and maximum recoil energies within a characteristic active-site volume. These physical fields constitute the sole quantitative input to the molecular layer.

### II.4. Molecular Layer – Gate Condition and Enzyme Disruption

#### Gate-condition statement

^28^Mg is chemically identical to stable Mg^2+^ prior to decay (identical atomic number, charge state and ionic radius). Thermodynamic binding affinities (*K_d_*) are therefore expected to be indistinguishable from those of native Mg^2+^ within ordinary experimental uncertainty. Secondary mass-dependent isotope effects on binding are generally small for light metals, and magnetic isotope effects are relevant primarily to the odd-mass, spin-½ nucleus ^25^Mg rather than the spin-zero nucleus ^28^Mg [9,10]. The framework consequently adopts the chemical-identity approximation

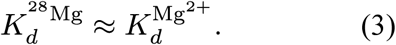

Fractional occupancy of any given magnesium-dependent active site is described by a competitive binding isotherm of hyperbolic (Michaelis–Menten-like) form [4,5]. Absolute free-energy calculations of binding lie outside the scope of the present study and remain an experimental prerequisite.

Within the computational framework, non-zero occupancy (*θ*_28_ > 0) is treated as an explicit input premise—the gate condition—with numerical values assigned explicit uncertainty bounds. All higher-scale predictions exist only under the assumption that this gate condition is satisfied. Once occupancy is assumed, nuclear transformation inside the active site generates two concurrent events: (i) abrupt change in cofactor charge and radius (Mg^2+^ → Al^3+^ → Si⁴⁺) and (ii) localized energy deposition. These events are converted into the Enzyme Disruption Index

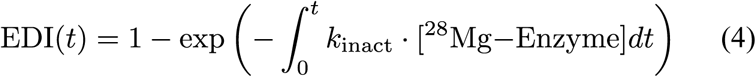

where *k_inact_* incorporates residence time and the probability of decay while the isotope occupies the site. EDI is computed separately for metabolic and DNA-repair enzyme classes and is interpreted as the fractional loss of catalytic capacity (range 0–1). The resulting catalytic-capacity modifiers are passed to the organelle layer.

### II.5. Organelle Layer

Molecular modifiers are aggregated into two organelle-scale state variables:

- Nuclear functional capacity (DNA-replication and repair throughput),
- Mitochondrial bioenergetic capacity (relative ATP-synthesis rate).

Each variable is a weighted average of the relevant EDI values, with weights derived from pathway annotation. The organelle capacities (values between 0 and 1) serve as direct inputs to the cellular layer.

### II.6. Cellular Layer

Organelle capacities are combined with the intrinsic difference in magnesium demand between malignant and normal cells to produce cellular fitness metrics (relative proliferation rate and metabolic survival probability). The replication-time asymmetry

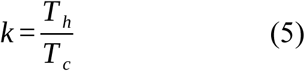

quantifies the proliferative advantage of tumor cells. Cellular fitness is expressed as a multiplicative factor that modulates the baseline tumor-cell growth rate and is transferred, together with *k*, to the tissue layer.

### II.7. Tissue Layer

Cellular fitness and replication asymmetry enter an intrinsic Gompertz formulation [11] in which the empirical deceleration parameter is replaced by biologically interpretable quantities:

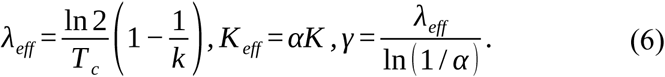

#### A critical transition threshold

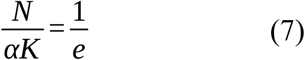

defines the boundary between an early, resource-limited growth regime and a progressively invasive regime. The interval between early detectability and this threshold constitutes the Intrinsic Therapeutic Window (ITW). Tumor-burden trajectories *N* (*t*) generated at this scale become the primary input to whole-body and clinical-prediction layers.

### II.8. Whole-body Layer

Systemic behavior is described by a hybrid physiologically based pharmacokinetic (PBPK) model adapted from ICRP Reference Man [12], with tumor-specific uptake driven by differential magnesium demand. Absorbed doses are calculated with the MIRD formalism [13] and corroborated by three-dimensional voxel Monte-Carlo transport. Differential tumor-versus-normal uptake fractions are treated as phenomenological parameters carrying explicit uncertainty (±50 %). Bone-marrow and whole-body doses are evaluated against QUANTEC constraints [14].

### II.9. Clinical Prediction Layer

Outputs from all preceding scales are integrated to estimate the Intrinsic Therapeutic Window and related timing metrics. These quantities are hypothesis-generating computational predictions; they do not constitute claims of clinical efficacy.

### II.10. Verification, Validation, and Uncertainty

Numerical stability, atom conservation, and consistency with NNDC, NIST E-STAR [15], and MIRD reference data were verified. Parameter-sensitivity and Monte-Carlo uncertainty analyses were performed on key biological and physical inputs. Cross-scale consistency was confirmed by verifying that monotonic changes in lower-scale parameters produce directionally coherent changes in higher-scale outputs. All biological parameters that lack direct experimental measurement for ^28^Mg are flagged as exploratory and are assigned explicit uncertainty ranges.

#### Code and Data Availability

The complete computational workflow, validation scripts, and supporting datasets will be deposited in a public repository upon publication. Supplementary Information contains the full parameter tables, audit logs, and Executive Report.

## III. Results

Assuming satisfaction of the gate condition—defined as non-zero fractional occupancy (*θ*_28_ > 0) of Mg^2+^-dependent catalytic sites by ^28^Mg, as governed by competitive binding isotherms of hyperbolic form [4,5]—the multiscale computational framework generated a sequence of emergent behaviors spanning six organizational levels, from atomic nuclear transformations to system-level therapeutic functions.

Nuclear decay data for the chain ^28^Mg → ^28^Al → ^28^Si (half-life of ^28^Mg = 20.915 h) were taken from the evaluated nuclear databases NNDC/ENSDF [8]. Results are organized according to the hierarchical propagation of mechanistic information defined in the Methods. At each organizational level, outputs generated by the preceding layer become constrained inputs to the next, allowing progressively higher-order biological behaviors to emerge. Throughout this study, all predictions remain strictly conditional upon successful catalytic-site occupancy and therefore represent computational consequences of the Radio-Cofactor Hypothesis rather than experimentally validated biological events.

### III.1. Emergence of the Radio-Cofactor Mechanism and the Enzyme Disruption Index

The first emergent behavior predicted by the framework arises immediately after the gate condition is assumed. Prior to catalytic occupancy, radionuclide decay is computationally indistinguishable from conventional intracellular irradiation. Once occupancy is assumed, radioactive decay becomes spatially confined to the enzyme–cofactor interface, converting the active site itself into the origin of coupled biochemical and radiophysical perturbations.

The molecular layer represented 312 Mg^2+^-dependent enzymes participating in DNA replication and repair, RNA transcription, ATP utilization, mitochondrial bioenergetics, cytoskeletal regulation, signal transduction, and intermediary metabolism. Four representative high-priority targets were retained to illustrate the computational workflow (Table 1). Their modeled catalytic inactivation-rate constants ranged from 0.0500 to 0.1000 h⁻¹.

**Table 1.**
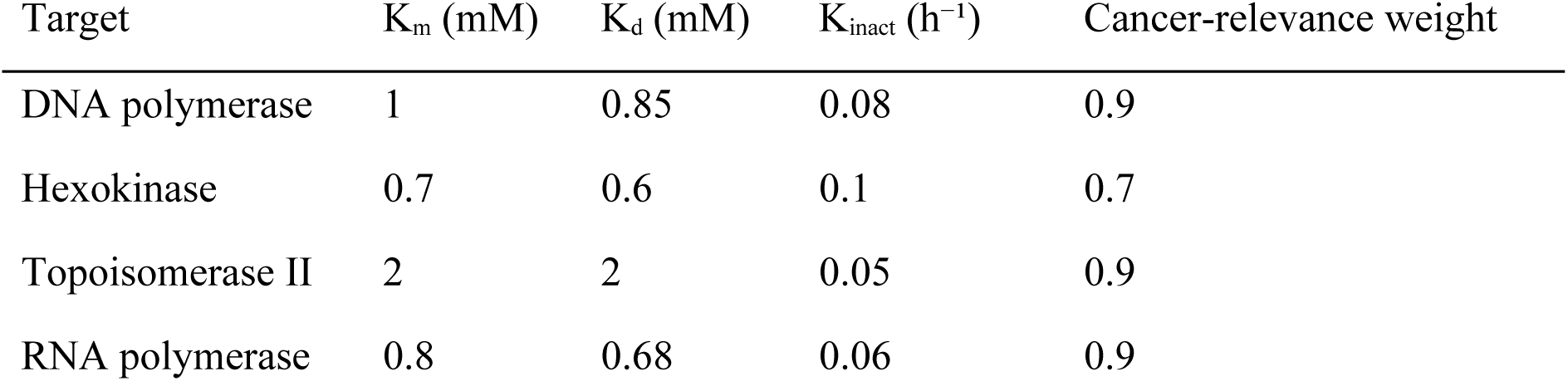
Representative Mg^2+^-dependent targets in the v10.2 enzyme module.

Following catalytic-site occupancy, the sequential decay chain ^28^Mg → ^28^Al → ^28^Si was propagated analytically with the Bateman equations (Figure 1). Nuclear kinetics reproduced the expected physical behavior with excellent numerical stability (maximum atom-conservation discrepancy 1.2207 × 10^-4^ atoms for an initial population of 10^12^ parent atoms). At one physical half-life of ^28^Mg (20.915 h), the model predicted 5.0008 × 10^11^ remaining parent atoms together with transient accumulation of 8.9623 × 10⁸ daughter ^28^Al atoms; after five half-lives more than 96 % of the initial inventory had converted to stable ^28^Si.

**Figure 1.**
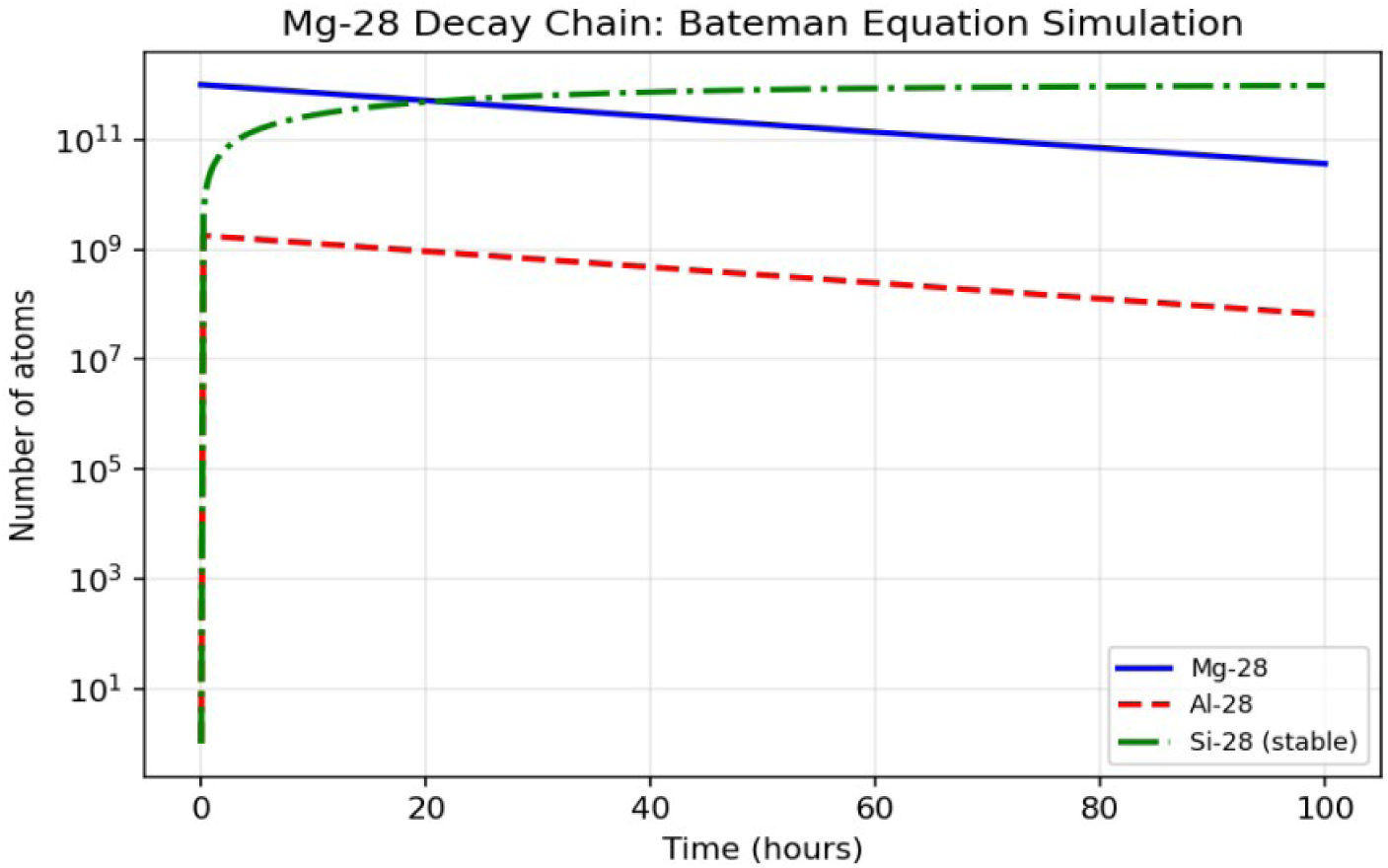
Bateman decay chain of ^28^Mg → ^28^Al → ^28^Si. Analytical solutions of the Bateman equations generated by Engine v10.2 for an initial population of 10^12^ ²□Mg atoms. The parent ²□Mg (solid blue), transient daughter ²□Al (dashed red), and stable ²□Si (dashed green) populations are shown on a logarithmic scale over 100 hours. At one half-life (20.915 h) the model predicts 5.0008 × 10^11^ parent atoms and 8.9623 × 10□ ²□Al atoms; after five half-lives the majority of the inventory has converted to stable ²□Si. Particle conservation is maintained (maximum numerical discrepancy 1.2207 × 10□□ atoms).

Decay occurring inside the catalytic pocket generates a mechanistically distinct event termed the Atomic Switch: simultaneous alteration of cofactor physicochemical identity (Mg^2+^ → Al^3+^ → Si⁴⁺) and deposition of localized radiation energy (Auger electrons, β-particles, and daughter-nucleus recoil) within nanometer dimensions of the active site (Figure 2). The two classes of output remain intrinsically coupled.

**Figure 2.**
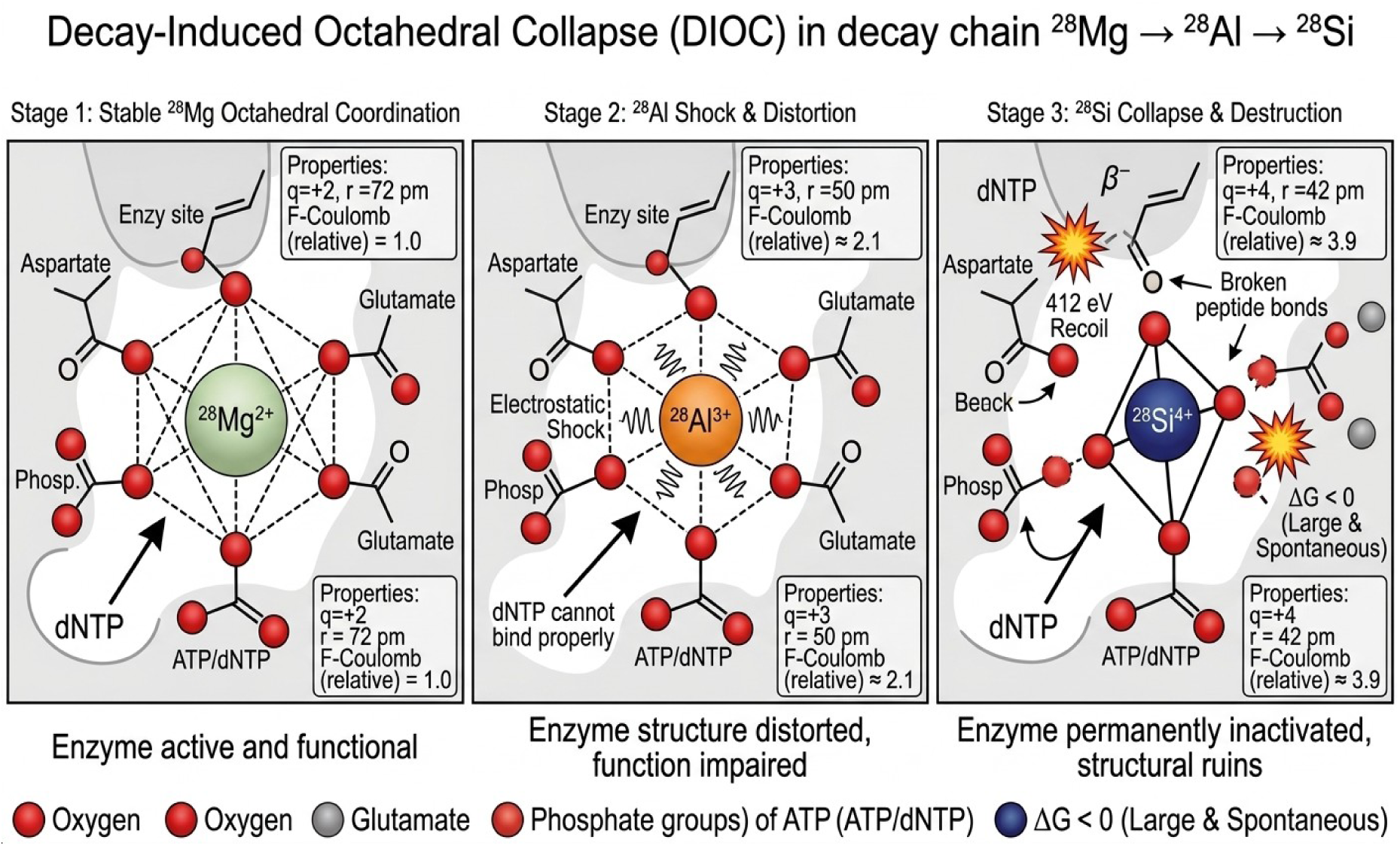
Enzyme Disruption Index (EDI) heatmap across 16 functional enzyme groups. Time-dependent EDI values predicted by the multiscale framework under the gate-condition assumption (Engine v10.2). Each row represents one functional enzyme class; columns correspond to 24 h, 48 h, 72 h and 96 h after administration of the reference single-cycle protocol. EDI is a deterministic model output quantifying fractional loss of catalytic capacity and has not been experimentally calibrated for ^28^Mg.

The Enzyme Disruption Index (EDI) isinterpreted as the fractional loss of catalytic capacity across the modeled enzyme network. Under the reference administration protocol, EDI increased progressively. In the representative U87MG glioblastoma parameterization predicted values rose from approximately 0.55–0.68 at 24 h to 0.92–0.96 at 48 h across major functional classes, reaching near-complete network disruption (≥0.99) by 72–96 h (Table 2 and Figure 3). Oxidative-phosphorylation and DNA-replication/repair enzymes showed among the highest early disruption. These results indicate that extensive impairment of intracellular catalytic networks is predicted to precede accumulation of high macroscopic absorbed tumor doses. Catalytic impairment was quantified by the Enzyme Disruption Index (Eq.4)

**Figure 3.**
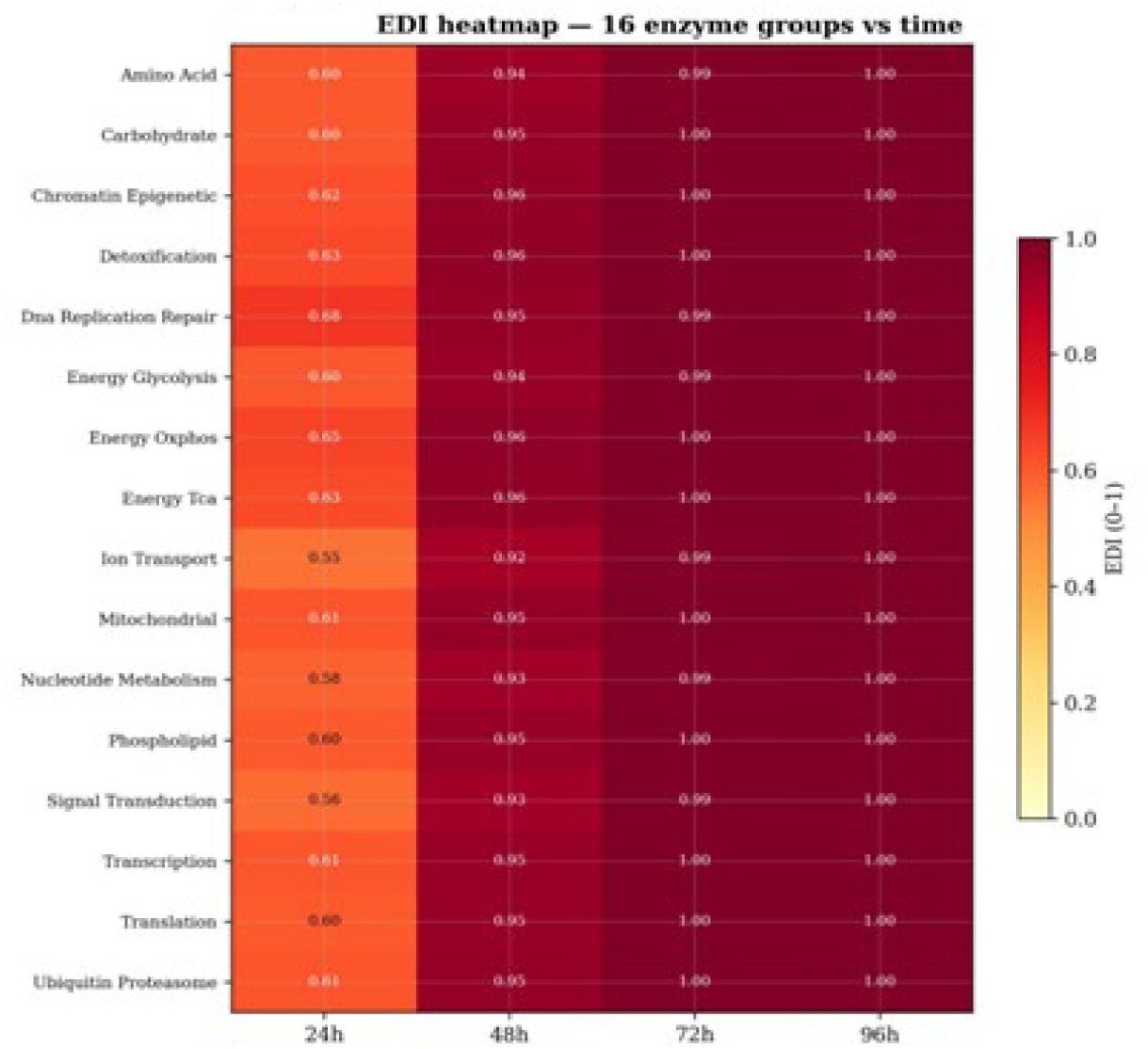
Decay-Induced Octahedral Collapse (DIOC) at a Mg^2+^-dependent catalytic site. Three-stage structural and electrostatic progression following nuclear transformation of ²□Mg that has occupied an enzyme active site (gate-condition assumption). Stage 1: Stable octahedral coordination of ²□Mg²□ (q = +2, r ≈ 72 pm) – enzyme remains functional. Stage 2: Sudden charge and radius change upon formation of ²□Al³□ (q = +3, r ≈ 50 pm) produces electrostatic shock and distortion of the coordination sphere. Stage 3: Further conversion to ²□Si□□ (q = +4, r ≈ 42 pm) together with 414 eV nuclear recoil leads to irreversible collapse of the active-site geometry, rupture of coordinating interactions, and permanent loss of catalytic function. The diagram is a schematic representation of the Atomic Switch predicted by Engine v10.2; it does not depict the preceding competitive occupancy step.

**Table 2.**
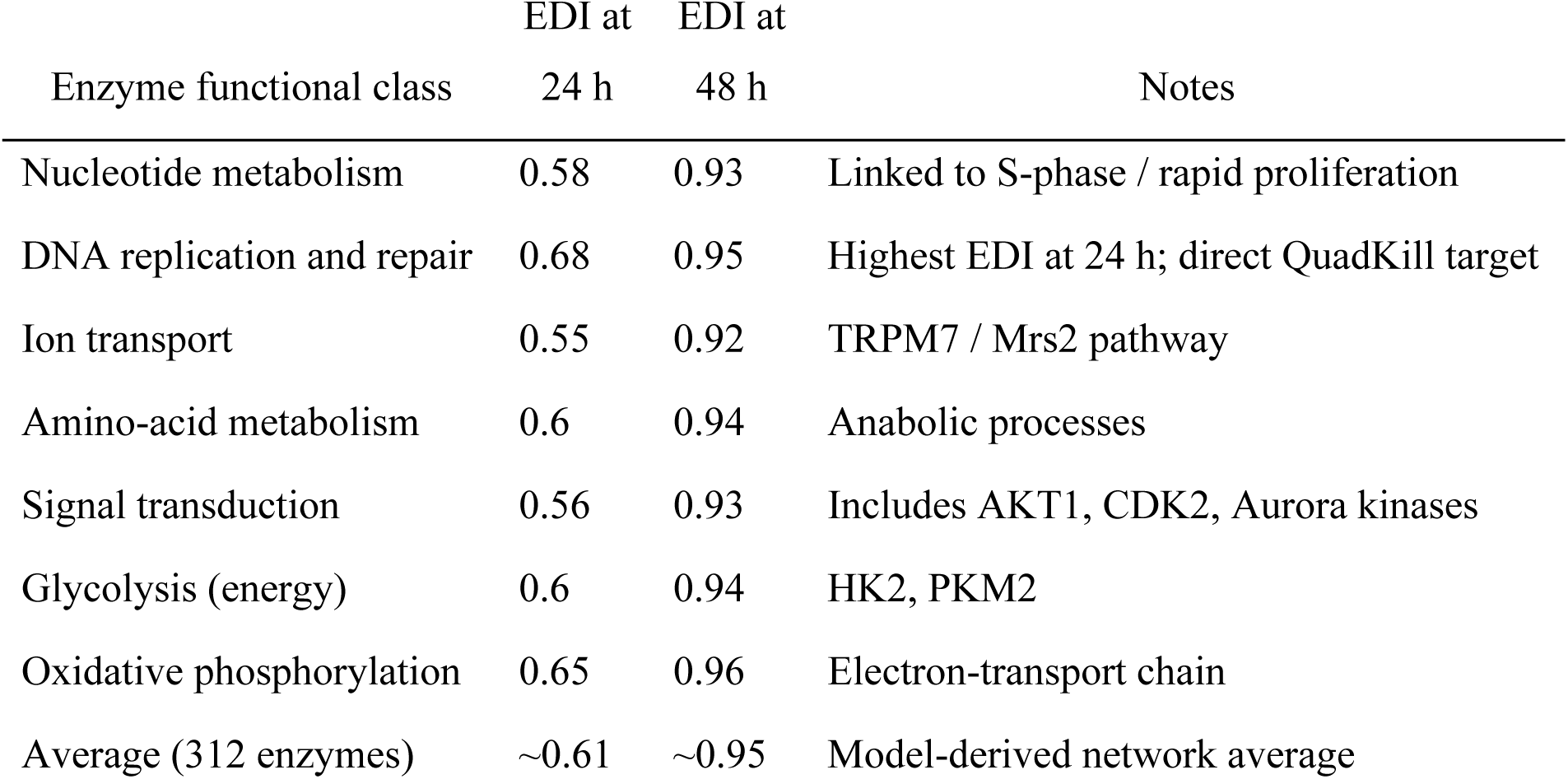
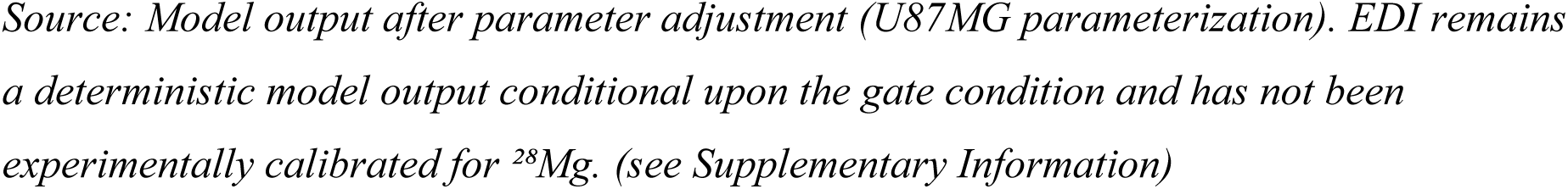
Enzyme Disruption Index (EDI) by functional class at 24 h and 48 h (U87MG parameterization).

| Enzyme functional class | EDI at 24 h | EDI at 48 h | Notes |
| --- | --- | --- | --- |
| Nucleotide metabolism | 0.58 | 0.93 | Linked to S-phase / rapid proliferation |
| DNA replication and repair | 0.68 | 0.95 | Highest EDI at 24 h; direct QuadKill target |
| Ion transport | 0.55 | 0.92 | TRPM7 / Mrs2 pathway |
| Amino-acid metabolism | 0.6 | 0.94 | Anabolic processes |
| Signal transduction | 0.56 | 0.93 | Includes AKT1, CDK2, Aurora kinases |
| Glycolysis (energy) | 0.6 | 0.94 | HK2, PKM2 |
| Oxidative phosphorylation | 0.65 | 0.96 | Electron-transport chain |
| Average (312 enzymes) | ~0.61 | ~0.95 | Model-derived network average |

*Source: Model output after parameter adjustment (U87MG parameterization). EDI remains a deterministic model output conditional upon the gate condition and has not been experimentally calibrated for $^{28}\text{Mg}$ . (see Supplementary Information)*

### III.2. Nuclear-decay parameters supporting the Atomic Switch

Nuclear and radiation-interaction parameters used throughout the framework are listed in Table 3. Mean β energies of 0.19 MeV (^28^Mg) and 1.50 MeV (^28^Al), the principal 1778.9-keV γ line of ^28^Al, the 1.39-keV Al KLL Auger component, and the 414-eV ^28^Si recoil energy were carried forward as localized perturbation inputs to the molecular layer. These physical quantities do not by themselves demonstrate biological effect; they supply the quantitative foundation for the EDI and subsequent organelle-scale responses.

**Table 3.**
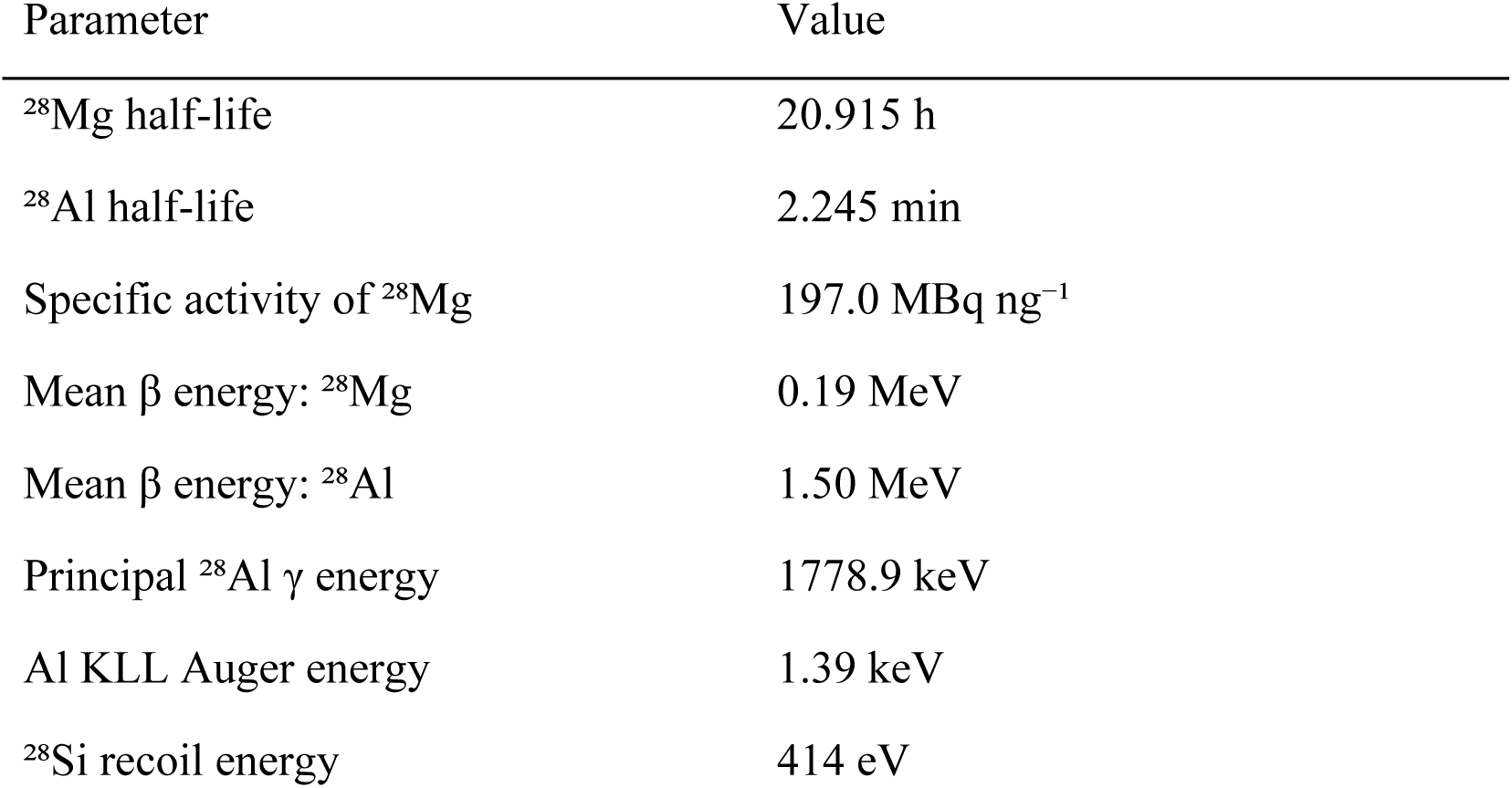
Nuclear and radiation-interaction parameters (v10.2).

### III.3. Propagation to organelle and cellular scales: the Quadruple-Kill cascade

Molecular-level catalytic-capacity modifiers were aggregated into two organelle-scale state variables: nuclear functional capacity (DNA-replication and repair throughput) and mitochondrial bioenergetic capacity (relative ATP-synthesis rate). Progressive loss of both capacities produced a coordinated reduction in cellular fitness (relative proliferation rate and metabolic survival probability).

The framework identifies this convergence as the Quadruple-Kill cascade—four mechanistically complementary injury pathways originating from a single radio-cofactor decay event (Figure 4 and Table 4): (i) irreversible disruption of Mg^2+^-dependent catalysis via the Atomic Switch, (ii) nanoscale energy deposition by Auger electrons, (iii) intracellular β-particle irradiation, and (iv) localized daughter-nucleus recoil. Because all four pathways arise from the same catalytic-site event, they evolve synchronously. Preferential amplification inside malignant cells is predicted to arise from the higher intrinsic magnesium demand of tumor cells rather than from receptor-mediated targeting.

**Figure 4.**
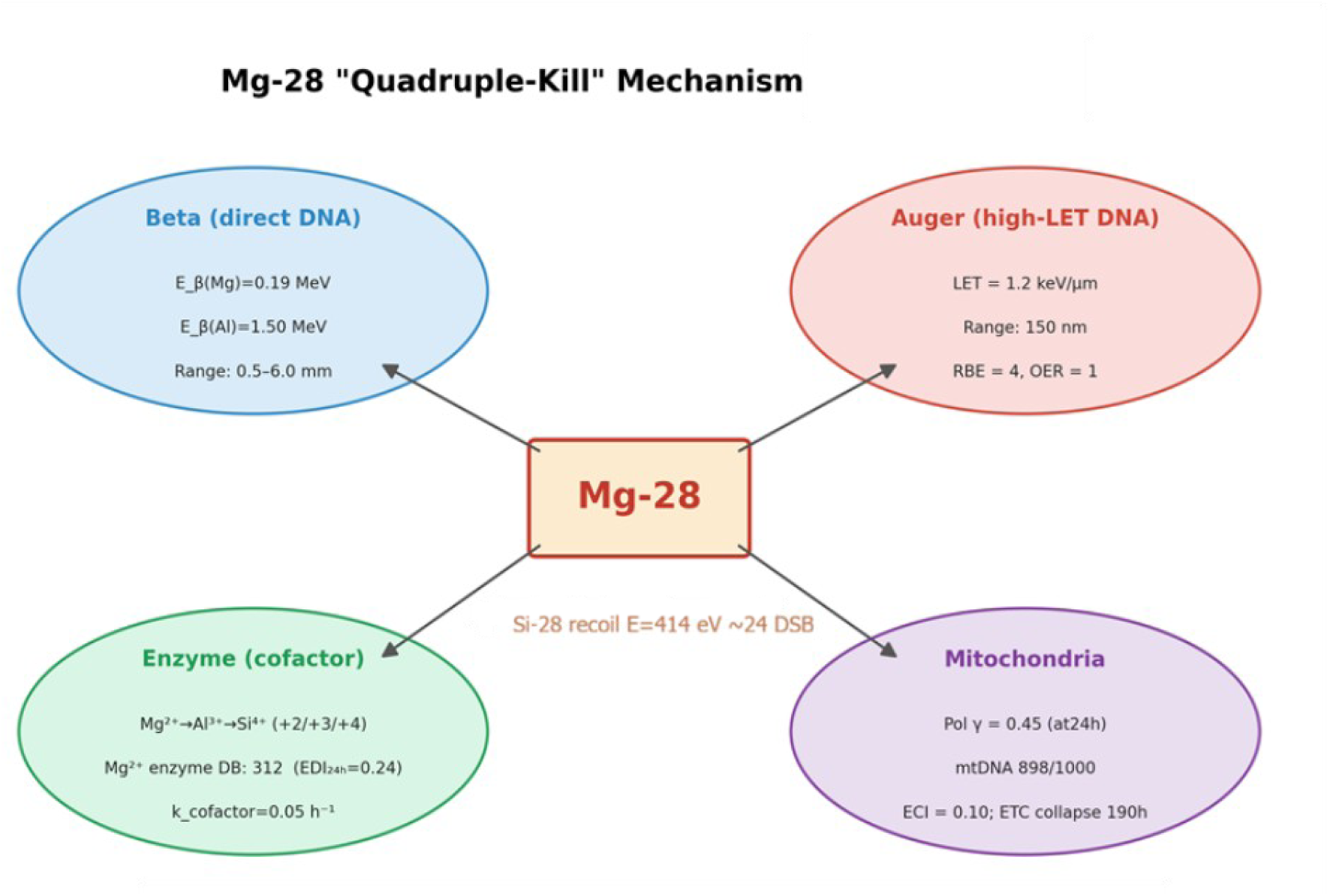
Schematic of the Quadruple-Kill cascade generated by a single ^28^Mg radio-cofactor decay event. The Atomic Switch (Mg^2+^ → Al^3+^ → Si^4+^) occurring inside an occupied catalytic site simultaneously initiates four coupled injury pathways: (i) irreversible loss of Mg²□-depenent enzymatic catalysis, (ii) high-LET Auger-electron deposition, (iii) intracellular β-particle irradiation, and (iv) localized ²□Si nuclear recoil (414 eV). The diagram illustrates the mechanistic convergence predicted by Engine v10.2; numerical values shown are model-derived and conditional upon satisfaction of the gate condition.

**Table 4.**
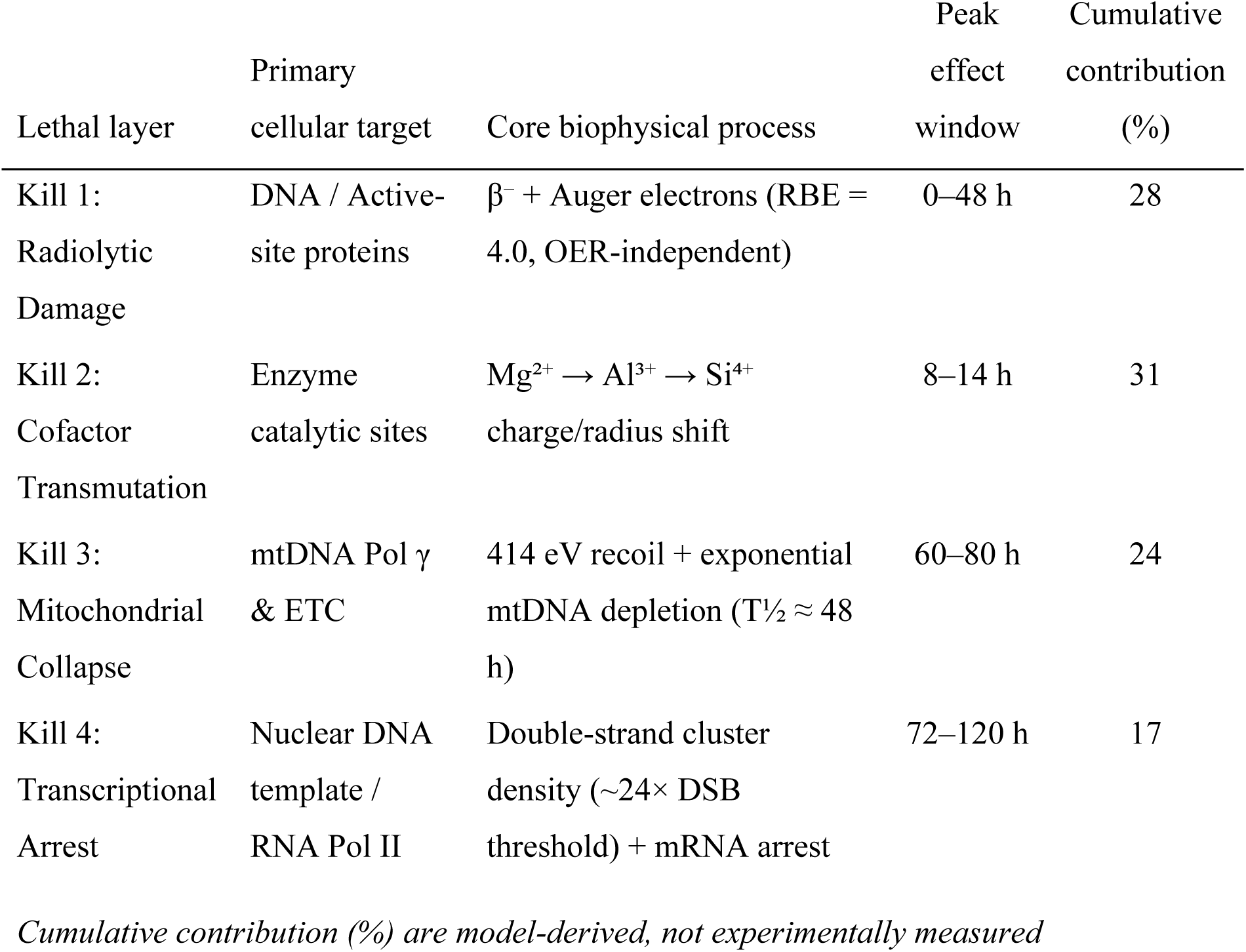
Quadruple-Kill layers and temporal contributions (Mean Over 90 Simulation Runs)

At the tissue scale the reduction in cellular fitness was inserted into the intrinsic Gompertz formulation, driving the normalized tumor burden N/(αK) toward values below the classical transition threshold of 1/e. Within the present study this trajectory is reported only as a qualitative systems-level illustration linking intracellular disruption to macroscopic tumor dynamics.

### III.4. Differential uptake and intrinsic imaging performance

Preferential magnesium uptake was encoded as relative coefficients referenced to normal cells: 3.5 (U87MG), 3.0 (PANC1), 2.8 (A549), 2.2 (HeLa), and 2.0 (MCF7) versus 1.0 for the normal-cell reference. These literature-informed coefficients define the self-targeting assumption used in the biodistribution and cellular-fitness modules; they are not direct measurements of ^28^Mg uptake.

The imaging module identified four γ peaks at 400.7, 941.7, 1342.3, and 1778.9 keV. Modeled scatter-to-primary ratios were 0.2244, 0.0760, 0.0500, and 0.0500, respectively. The 1778.9-keV ^28^Al line was selected as the preferred intrinsic imaging signal (simulated blur 0.50 mm, SPR 0.0500), generating a single-isotope imaging hypothesis that can be tested in prospective detector and biodistribution studies (Figure 5).

**Figure 5.**
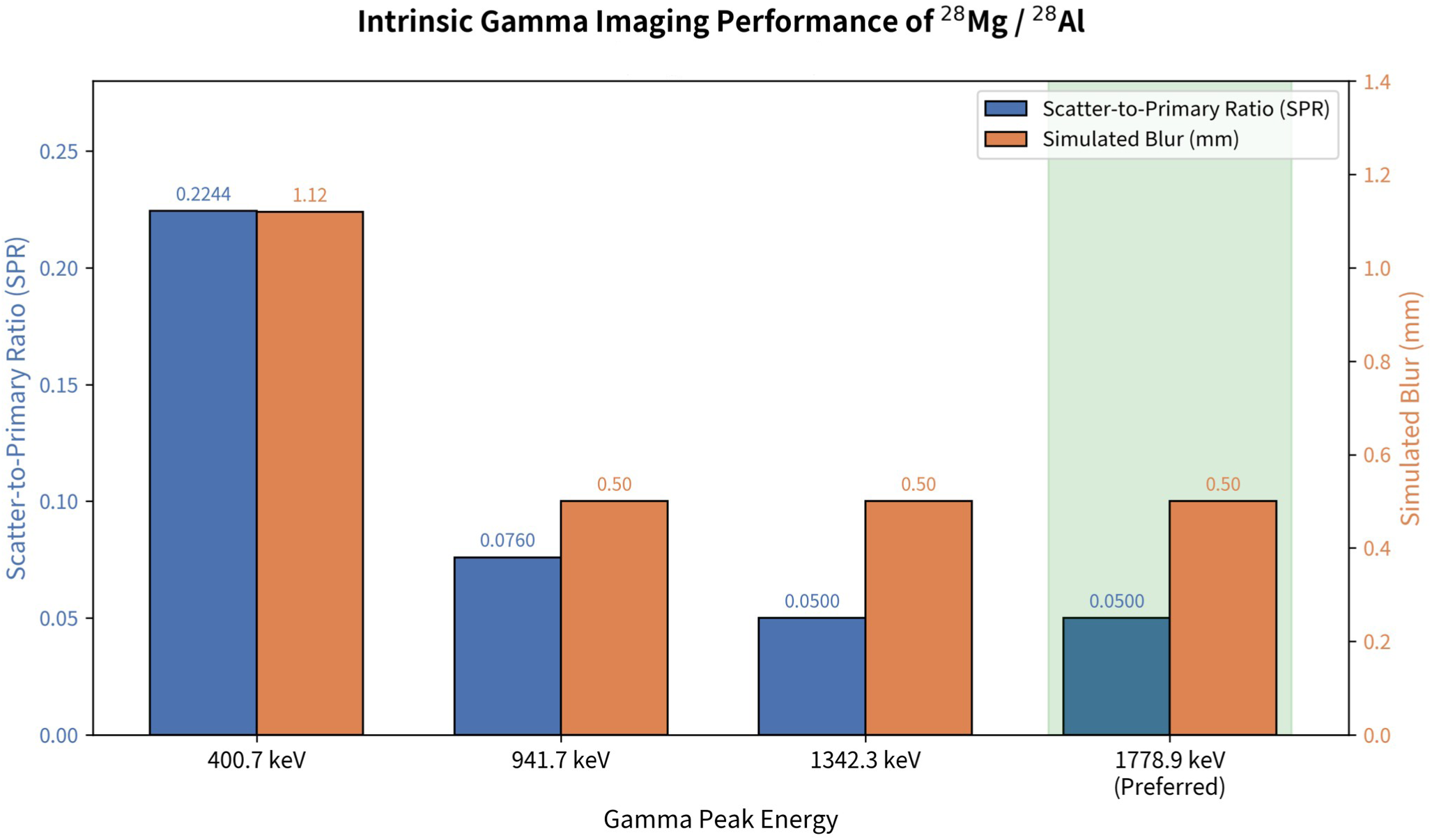
Intrinsic gamma imaging performance of ^2+^Mg / ^28^Al. Scatter-to-primary ratio (SPR, blue) and simulated spatial blur (orange) for the four principal gamma peaks identified by the imaging module (Engine v10.2). The 1778.9 keV ² □Al line (highlighted) exhibits the lowest SPR (0.0500) and the smallest blur (0.50 mm), and is therefore selected as the preferred intrinsic imaging signal.

### III.5. Pharmacokinetics, dosimetry, and QUANTEC-constrained safety screen

In the two-compartment intravenous-bolus model, blood and tissue retention times were 7.1922 h and 106.7461 h. Extension to a three-compartment model incorporating tumor yielded a peak modeled tumor activity of 0.3729 MBq at 9.63 h and a tumor time-integrated activity of 21.1819 MBq·h. Biodistribution fractions (tumor 5 %, muscle 30 %, bone 25 %, kidneys 15 %, liver 10 %, gastrointestinal tract 8 %, blood 5 %) carry explicit uncertainty of approximately ±50 % because direct in-vivo ^28^Mg data are unavailable.

For the safety-constrained single-cycle reference scenario (0.3 ng = 59.1 MBq; 5-g reference tumor) the MIRD module estimated absorbed doses of 1.636 Gy (bone), 0.931 Gy (kidneys), 0.817 Gy (liver), 0.614 Gy (tumor), and 0.327 Gy (whole body) (Table 5 and Figure 6). Modeled bone and whole-body values lie below the corresponding QUANTEC 2010 reference limits of 2.0 Gy and 0.6 Gy. Calculated tumor-to-kidney and tumor-to-bone therapeutic indices were 3.278 and 1.864, respectively.

**Table 5.**
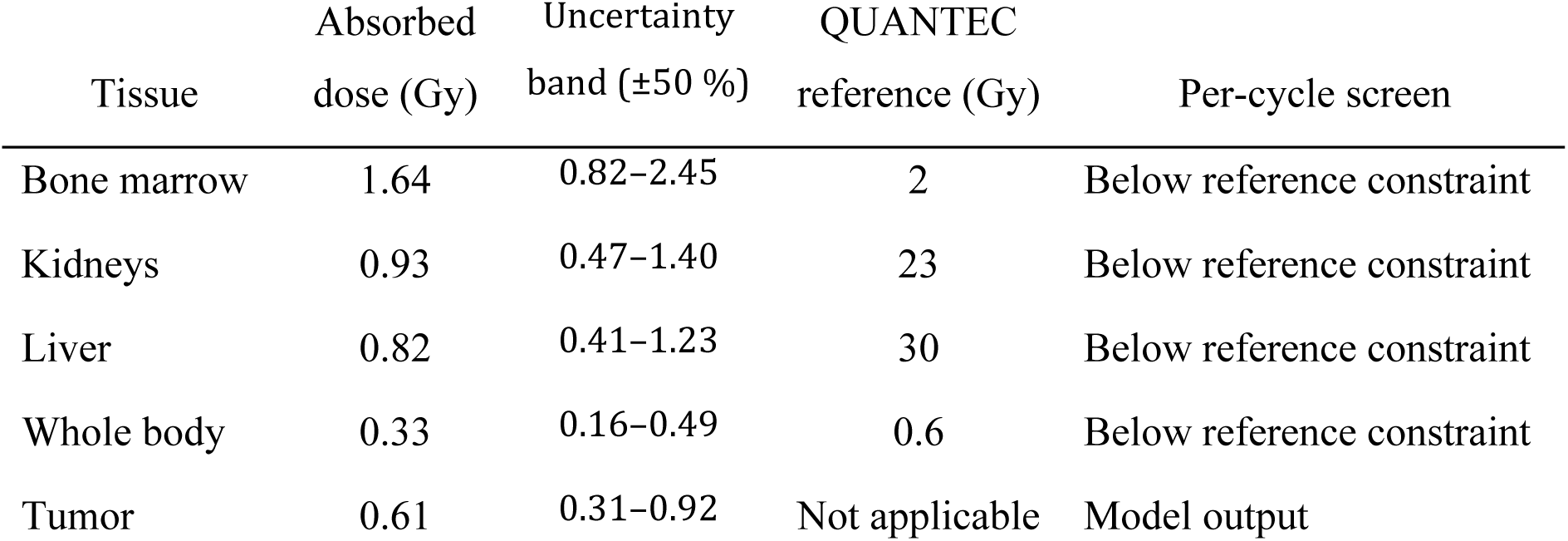
Modelled absorbed doses for the v10.2 single-cycle reference scenario (0.3 ng; 59.1 MBq; 5-g tumor).

**Figure 6.**
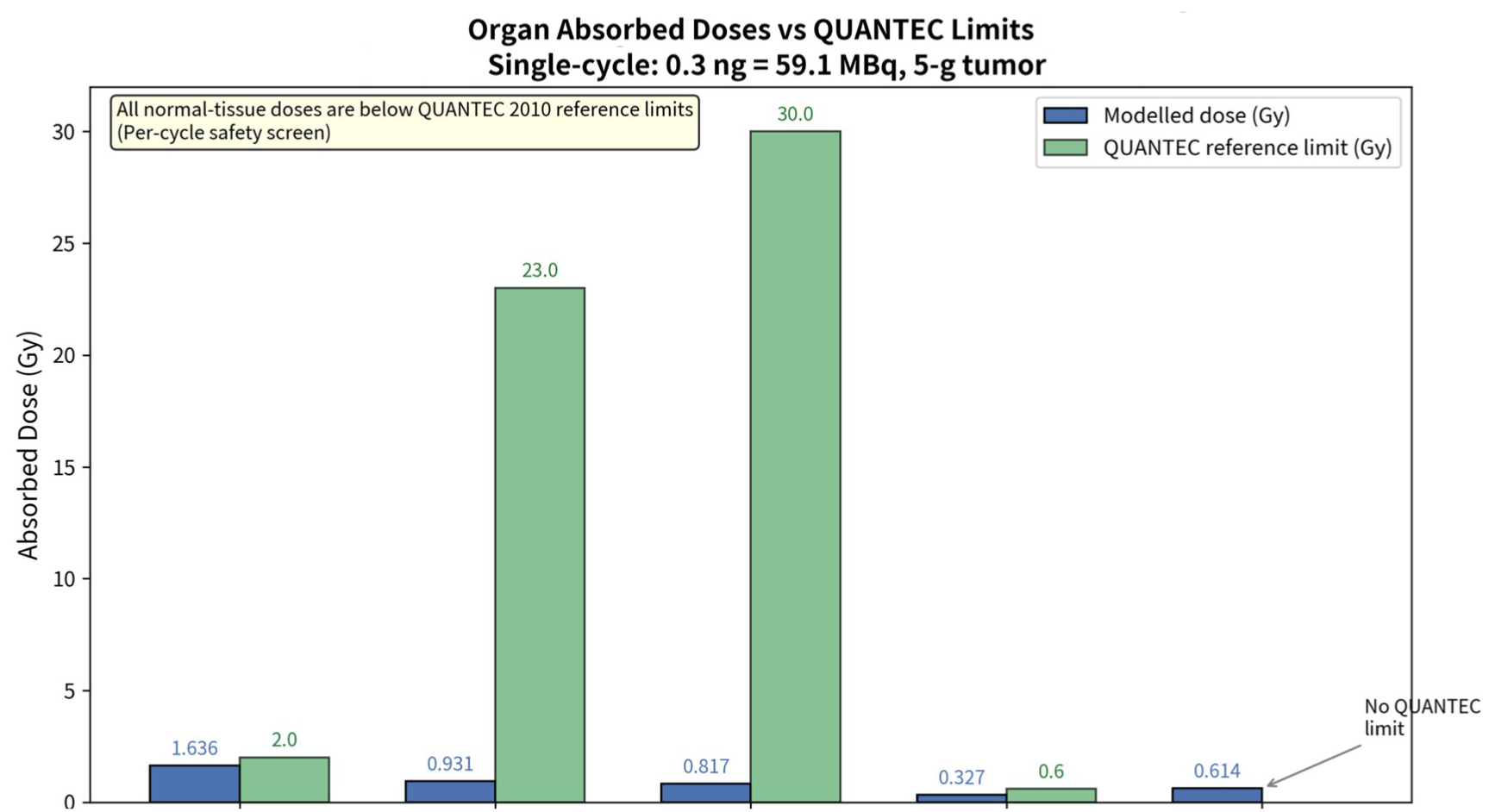
Organ absorbed doses versus QUANTEC 2010 reference limits. Modelled absorbed doses for the safety-constrained single-cycle reference scenario (0.3 ng = 59.1 MBq; 5-g reference tumor) calculated with the MIRD module of Engine v10.2. Blue bars show modelled doses; green bars show the corresponding QUANTEC reference limits. All normal-tissue doses lie below the reference constraints (per-cycle safety screen). Tumor dose is reported without a QUANTEC limit.

The calculated tumour absorbed dose of approximately 0.61 Gy obtained with the safety-constrained single-cycle activity of 59.1 MBq (0.3 ng) and a 5 g reference tumour is a direct consequence of two deliberate modelling choices: (i) the phenomenological tumour uptake fraction of only 5 % (adapted from ICRP Reference Man magnesium distribution and carrying an explicit uncertainty of ±50 % in the absence of direct in-vivo ^28^Mg biodistribution data), and (ii) the selection of a low administered activity designed to remain below the corresponding QUANTEC 2010 per-cycle constraints for bone marrow and whole body.

For physical reference, the theoretical self-dose under the assumption of 100 % tumour absorption would be substantially higher (approximately 1053 Gy for a 10 g lesion after six cycles of 0.3 ng). Because the primary therapeutic leverage proposed in this framework is the localised catalytic-site Atomic Switch together with Auger-electron and nuclear-recoil effects rather than macroscopic absorbed dose, the model prioritises normal-tissue safety. Should experimental biodistribution studies confirm a higher tumour uptake fraction, the tumour dose would scale proportionally while still permitting the same per-cycle safety screen. All normal-tissue doses reported in Table 5 remain below the corresponding QUANTEC limits at the point estimate; the upper bound of the ±50 % uncertainty band for bone marrow (2.45 Gy) exceeds the 2.0 Gy reference and therefore constitutes a residual risk that must be resolved by direct in-vivo biodistribution measurements.

The selected protocol is 0.3 ng per administration for six cycles separated by 72 h. The QUANTEC comparison is implemented strictly as a per-cycle safety screen; cumulative multi-cycle claims would require an explicit time-resolved marrow-recovery model followed by in-vivo toxicity validation.

### III.6. Comparative self-theranostic profile

In the model, ^28^Mg differs from comparator radionuclides by combining β emission with Auger-electron and recoil components, retaining intrinsic γ emissions for imaging, and being modeled without an exogenous ligand or carrier (Table 6). Relative efficiency values were calculated on a per-GBq model basis and must be interpreted strictly as comparative simulation outputs.

**Table 6.**
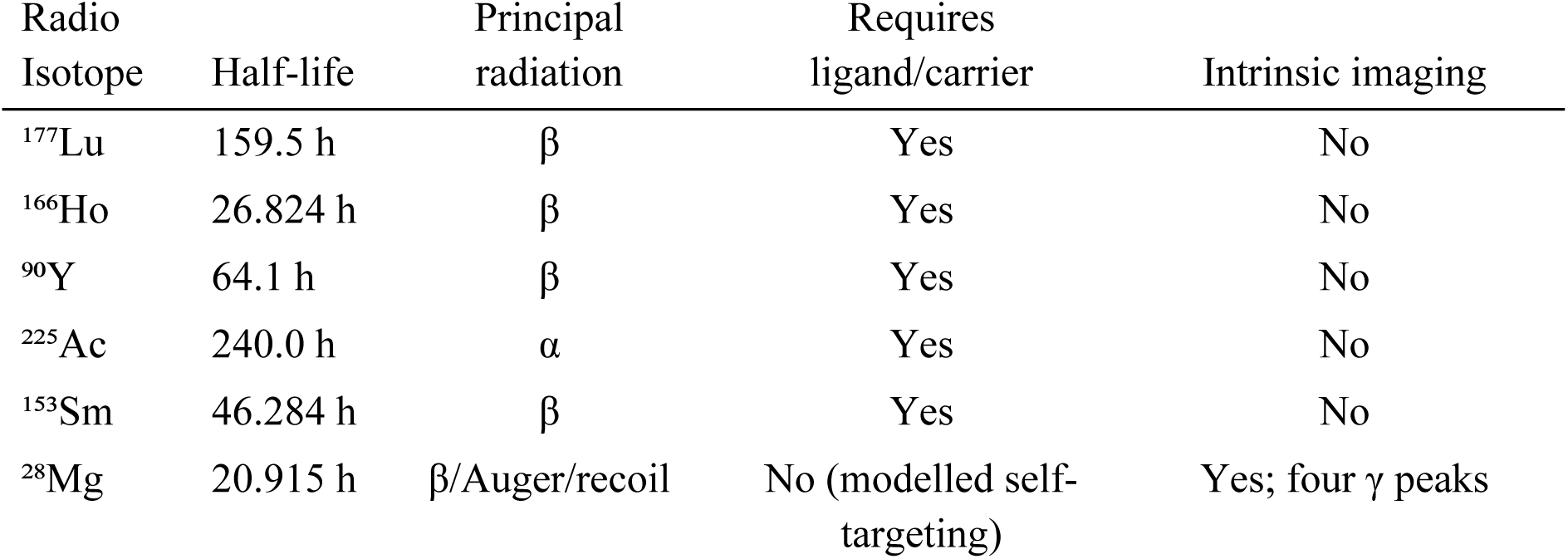
Comparison of ^28^Mg with selected therapeutic radionuclides.

### III.7. Evidence level and experimental transition

Nuclear constants and decay-chain calculations are supported by reference nuclear data (NNDC/ENSDF 2024). Enzyme inhibition, preferential uptake, biodistribution, organ doses, and therapeutic indices remain model-derived. The evidence matrix therefore classifies nuclear physics as high-confidence, the QuadKill mechanism and dosimetry as theoretically or computationally supported, and in-vivo efficacy as unestablished.

Preclinical priorities defined by the computational results are:

1. Cell-free enzyme assays that correlate EDI metrics with measured catalytic activity.
2. Confirmation of differential magnesium uptake in at least three cancer cell lines, performed first with isotopically enriched, non-radioactive ^25^Mg (the only stable magnesium isotope with nuclear spin I = 5/2). Uptake and intracellular distribution will be quantified by ICP-MS and complemented by ^25^Mg NMR spectroscopy to probe coordination environment. Only after these Phase-0 studies establish preferential uptake will analogous experiments proceed with ^28^Mg.
3. Single-dose xenograft biodistribution.
4. Multi-cycle toxicity assessment with hematologic monitoring.

In this way the computational outputs define measurable decision points for experimental validation rather than a claim of therapeutic readiness.

## IV. Discussion

### IV.1. Mechanistic interpretation of the Gate Condition

The Gate Condition introduced in the present framework can be further interpreted using a minimal kinetic formulation that links tumor growth dynamics with intracellular radio-cofactor availability. Rather than treating intracellular occupancy as an independent premise, the model considers the balance between radio-cofactor influx and radioactive decay.

The intracellular concentration of ^28^Mg is assumed to obey the mass-balance equation

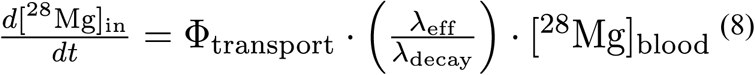

where *λ_eff_* is the effective resource-capture constant derived from tumor replication asymmetry, *λ_decay_* is the radioactive decay constant of ^28^Mg, and *Φ_transport_* (0–1) is an effective transport coefficient. The latter summarizes the combined influence of the principal magnesium transport pathways—primarily the plasma-membrane channels TRPM6/TRPM7 [16] and MagT1 [17], the mitochondrial uptake channel MRS2 [18], and the secondary contribution of efflux systems such as SLC41A1 and the CNNM family.

Under the quasi-steady-state approximation ((*d*[^28^Mg]_in_/*dt* ≈ 0, intracellular availability simplifies to

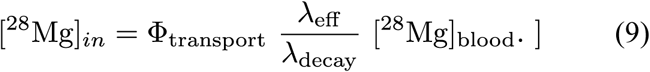

This expression is phenomenological rather than thermodynamic. It reflects the competition between two opposing processes: biological resource capture by proliferating tumor cells and radioactive loss through nuclear decay. Substituting the relationship into the competitive binding isotherm yields

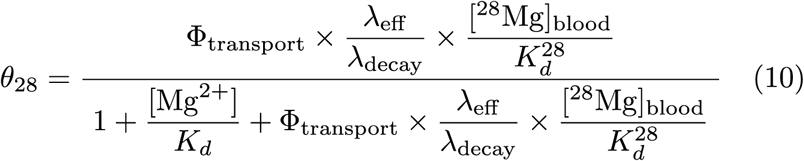

where *θ*_28_ denotes the fractional occupancy of magnesium-dependent catalytic sites by ^28^Mg. The formulation provides a mechanistic bridge between tumor growth kinetics and molecular targeting. Tumors characterized by greater replication asymmetry possess larger *λ_eff_* and, typically, elevated expression or activity of key influx transporters (TRPM7, MagT1). The resulting increase in effective intracellular availability of ^28^Mg raises the probability of catalytic-site occupancy. Conversely, when *λ_eff_ →* 0, intracellular availability approaches zero and *θ*_28_ becomes negligible, preventing initiation of the radio-cofactor mechanism.

Importantly, the model does not imply that tumor aggressiveness alters the intrinsic thermodynamic binding affinity (*K_d_*) of *Mg*^2+^. The chemical identity of magnesium prior to radioactive decay is assumed to remain unchanged. Instead, tumor growth dynamics and the associated transport machinery are hypothesized to regulate the effective intracellular concentration of ^28^Mg, thereby determining the degree to which the molecular gate is opened. Once this continuous (rather than binary) gate condition is satisfied, radioactive decay within the occupied catalytic site initiates the downstream cascade of enzyme disruption (EDI), organelle dysfunction, cellular failure, and ultimately tumor response.

This minimal kinetic extension is offered as a proposed interpretive framework rather than a mandatory premise of the computational results presented earlier. Its principal value lies in converting the Gate Condition from a qualitative switch into a quantitatively linked, experimentally approachable quantity that can, in principle, be tested by measuring differential transporter expression, *Mg*^2+^ uptake kinetics, and intracellular free-magnesium dynamics in matched tumor and normal cell populations. The kinetic interpretation developed above naturally extends beyond ^28^Mg and suggests a broader framework for evaluating other candidate radio-cofactors.

#### Extension of the Gate Condition to a Preliminary Radio-Cofactor Screening Framework

The intracellular availability of a candidate radio-cofactor is governed by the competition between biological resource capture and radioactive decay. Under the quasi-steady-state mass-balance relation introduced above, the first-order dynamic gate index is defined as

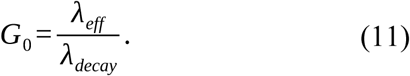

A higher *G*_0_ indicates a greater theoretical opportunity for intracellular accumulation before radioactive depletion, provided that suitable transporters and molecular compatibility are present. The Gate index itself does not measure therapeutic efficacy; it quantifies only the intrinsic kinetic competition between enrichment and decay.

Because radio-cofactor therapy requires simultaneous satisfaction of multiple biological and physical constraints, the Gate concept is extended into a preliminary screening matrix that incorporates transporter availability, cofactor compatibility, decay characteristics, organellar localization, enzyme-disruption potential, and normal-tissue dosimetric considerations (Table 7).

**Table 7.**
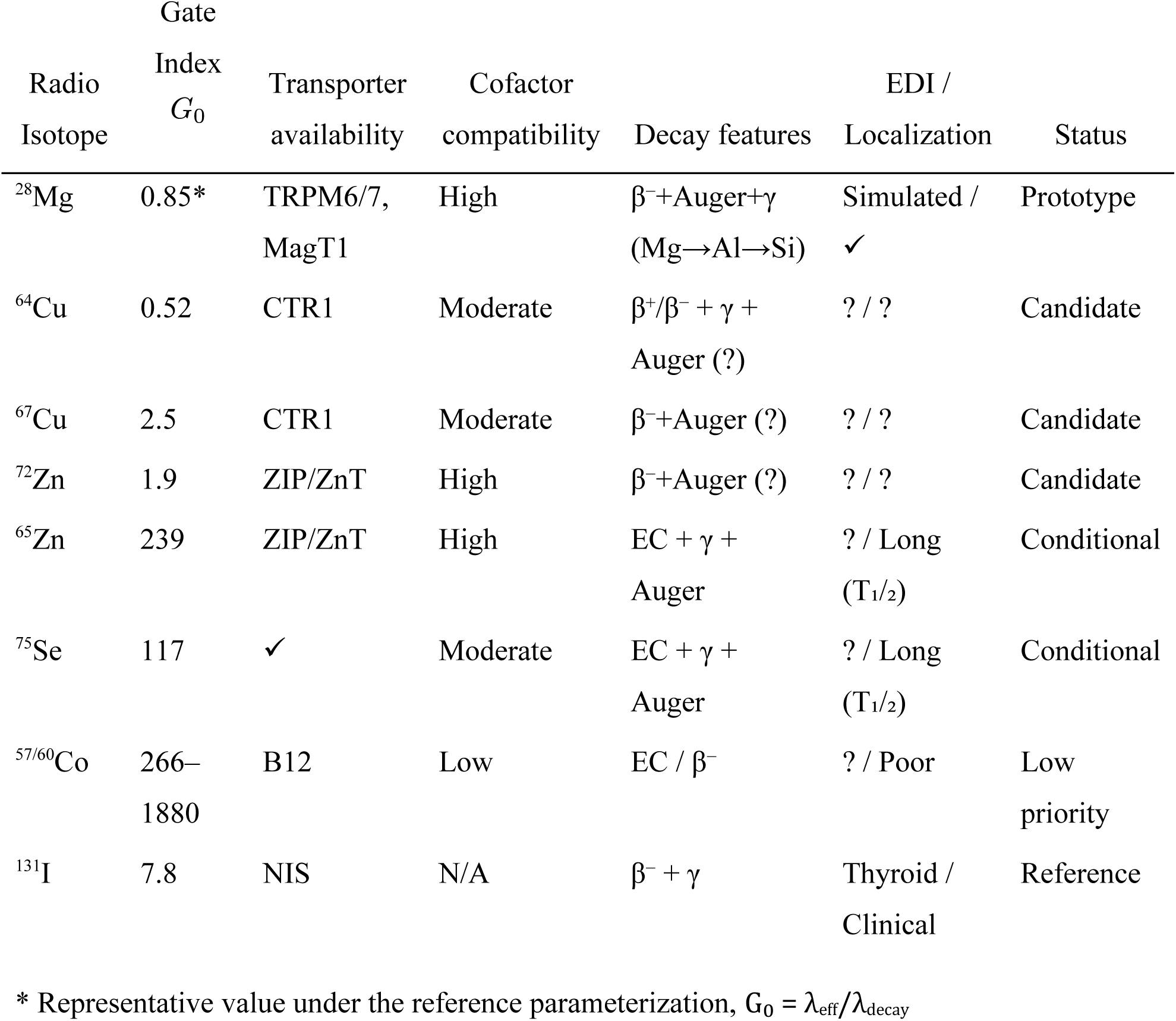
Preliminary computational screening matrix for candidate radio-cofactors derived from biologically essential metallic elements. Filled entries summarize currently available evidence; question marks denote research gaps. EDI has been quantified only for ^28^Mg.

The matrix is not intended as a definitive ranking. ^28^Mg remains the only fully evaluated prototype within the present framework. For this isotope the combination of high magnesium dependency in proliferating cells, established transport pathways (TRPM6/7, MagT1, MRS2), and the sequential nuclear transformation ^28^Mg → ^28^Al → ^28^Si creates a unique opportunity for coordinated chemical and radiological disruption. Comparative EDI values for the other candidates remain undetermined because isotope-specific localization and decay-response simulations have not yet been performed.

Decay characteristics further refine the screening. Auger electrons deposit energy on the nanometer scale and are therefore expected to be most effective when the radionuclide reaches a sensitive catalytic or nucleic-acid-associated compartment. Beta particles offer longer-range irradiation that may be advantageous in heterogeneous tumors, while gamma emissions can provide intrinsic imaging capability at the cost of additional normal-tissue exposure. These considerations reinforce the requirement that a viable radio-cofactor must not only satisfy the Gate condition but also achieve appropriate organellar localization.

Thus the screening framework converts the Gate Condition from a binary premise into a multi-dimensional filter that can guide future isotope-specific computational and experimental work while preserving ^28^Mg as the current prototype.

### IV.2. From occupancy to the Atomic Switch / DIOC and EDI

Once non-zero fractional occupancy (*θ*_28_ > 0) is achieved, the subsequent nuclear transformation of ^28^Mg within the catalytic pocket generates a mechanistically distinct event that the framework terms the Atomic Switch. The sequential decay ^28^Mg → ^28^Al → ^28^Si produces two concurrent perturbations: (i) an abrupt change in cofactor physicochemical identity (charge and ionic radius progressing from *Mg*^2+^ (≈72 pm) through *Al*^3+^ (≈50 pm) to *Si*^4+^ (≈42 pm)) and (ii) localized energy deposition by Auger electrons, *β*-particles and the 414 eV nuclear recoil of the daughter ^28^Si.

The structural and electrostatic consequences of this dual perturbation are illustrated schematically as Decay-Induced Octahedral Collapse (DIOC). In the first stage the octahedral coordination geometry remains intact and catalytic function is preserved. Upon formation of the ^28^*Al*^3+^ sudden increase in charge density and contraction of ionic radius generate an electrostatic shock that distorts the ligand field. Further conversion to ^28^*Si*^4^ together with the recoil impulse irreversibly disrupts coordinating interactions, collapsing the active-site geometry and abolishing catalytic competence.

These molecular events are quantified by the Enzyme Disruption Index (Eq. 4). which represents the fractional loss of catalytic capacity across the modelled network of magnesium-dependent enzymes. Because EDI is computed separately for distinct functional classes (DNA replication/repair, oxidative phosphorylation, glycolysis, signal transduction, etc.), the framework captures differential temporal profiles of catalytic impairment. Under the reference parameterization, network-averaged EDI rises rapidly, reaching near-complete disruption within 72–96 h. Critically, this progressive loss of catalytic capacity precedes the accumulation of high macroscopic absorbed doses, indicating that the primary therapeutic leverage of the radio-cofactor concept operates at the molecular rather than the macroscopic dosimetric scale.

### IV.3. Emergence and coordination of the Quadruple-Kill cascade

Aggregation of the catalytic-capacity modifiers into organelle-scale state variables (nuclear functional capacity and mitochondrial bioenergetic capacity) produces a coordinated reduction in cellular fitness. The framework identifies this multi-level convergence as the Quadruple-Kill cascade—four mechanistically complementary injury pathways that originate from a single catalytic-site decay event:

1. irreversible loss of *Mg*^2+^-dependent enzymatic catalysis via the Atomic Switch / DIOC,
2. nanoscale high-LET energy deposition by Auger electrons,
3. intracellular *β*-particle irradiation, and
4. localized nuclear recoil of the daughter ^28^Si.

Because all four pathways are spatially and temporally coupled to the same occupancy event, they evolve synchronously rather than as independent stochastic processes. Preferential amplification inside malignant cells is predicted to arise from the higher intrinsic magnesium demand and elevated transporter activity of tumor cells, rather than from receptor-mediated targeting. At the tissue scale the resultant reduction in cellular fitness is inserted into the intrinsic Gompertz formulation, driving normalized tumor burden toward values below the classical transition threshold of 1 / *e*. The cascade therefore constitutes a computational hypothesis of coordinated multi-scale disruption whose validity remains contingent upon experimental confirmation of the upstream gate condition.

Importantly, the Gompertz trajectories shown in Figure 7 should not be interpreted as an independent therapeutic mechanism. Rather, they represent the systems-level manifestation of the multiscale framework. Once the Gate Condition is satisfied, coordinated intracellular disruption—through the Atomic Switch, progressive EDI, and the Quadruple-Kill cascade—reduces tumor cellular fitness, causing tumor burden to evolve below the classical Gompertz transition threshold (N/αK = 1/e). In this framework, macroscopic tumor regression emerges from coordinated multi-target intracellular disruption, whereas absorbed radiation dose constitutes one contributing component rather than the sole therapeutic mechanism.

**Figure 7.**
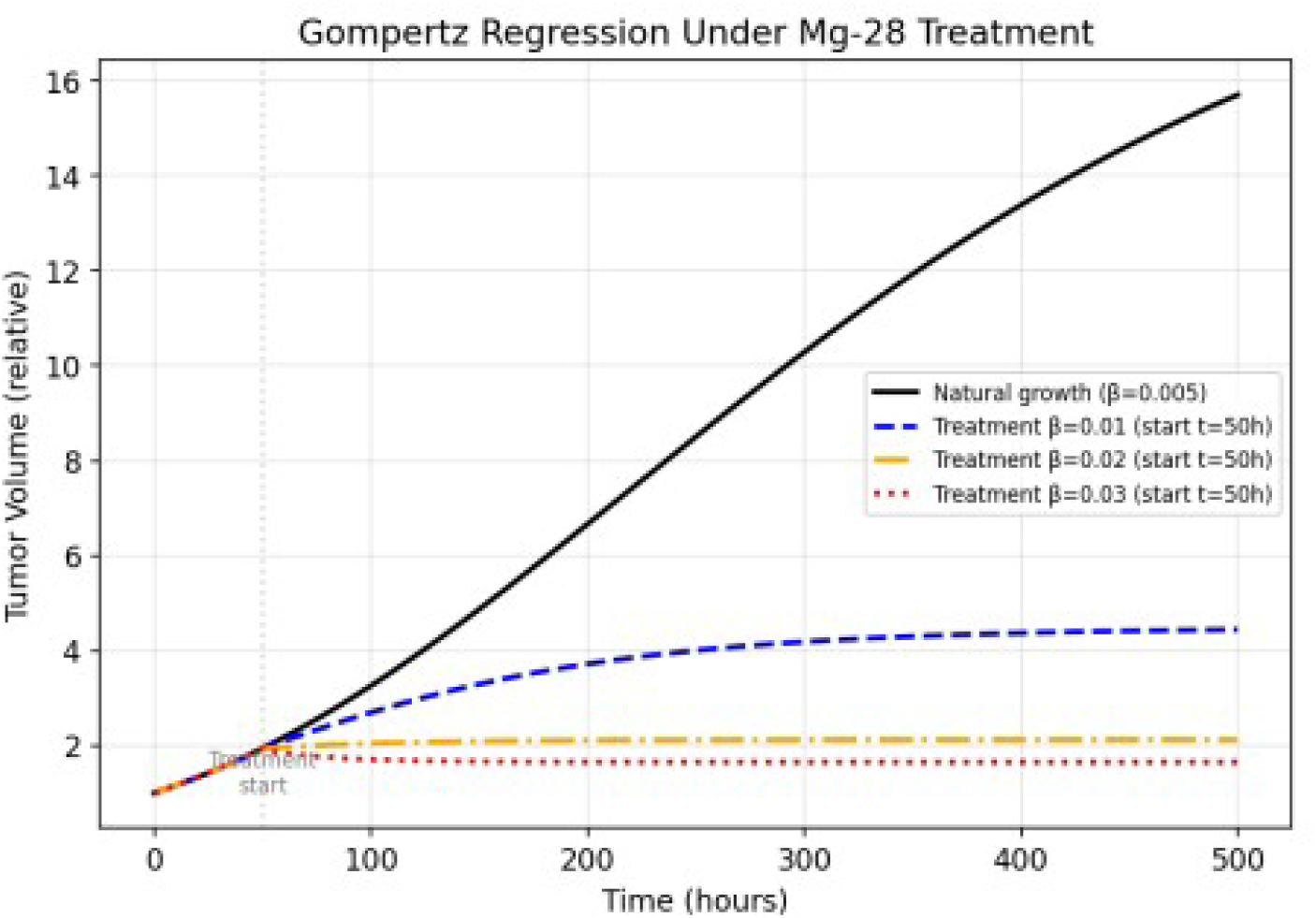
Illustrative Gompertz tumor-volume trajectories under the Radio-Cofactor mechanism. Relative tumor volume predicted by the intrinsic Gompertz formulation when cellular fitness is reduced by the Quadruple-Kill cascade (Engine v10.2). The solid black curve represents natural growth (β = 0.005). Dashed and dotted curves illustrate the effect of increasing effective kill rates (β = 0.01, 0.02 and 0.03) applied from t = 50 h. These trajectories are qualitative model illustrations generated under assumed parameter values; they are not experimentally validated tumor-volume reductions and should be interpreted strictly as exploratory computational projections.

### IV.4. System-level integration: the Quintax Functional Model

The hierarchical propagation of information from atomic decay through molecular occupancy, organelle dysfunction and cellular fitness culminates in a system-level construct that the framework designates the Quintax Functional Model. Quintax integrates five interdependent control axes—(i) catalytic-site occupancy, (ii) enzyme-network disruption, (iii) organelle-capacity collapse, (iv) differential cellular fitness, and (v) tissue-scale growth modulation—into a single coherent description of radio-cofactor activity.

Within this architecture each axis both receives constrained inputs from the level below and imposes top-down constraints on the level above. The model therefore embodies the dual bottom-up / top-down information flow formalized in the Methods. Importantly, Quintax is not offered as a predictive clinical algorithm; it is a hypothesis-generating scaffold that organizes the emergent behaviors identified by the multiscale simulations and that highlights the critical decision points at which experimental interrogation can falsify or refine the underlying assumptions.

### IV.5. Pharmacokinetics, dosimetry and QUANTEC-constrained safety

Systemic behavior is described by a hybrid physiologically based pharmacokinetic model with tumor-specific uptake driven by differential magnesium demand. Absorbed doses calculated with the MIRD formalism for the safety-constrained single-cycle reference scenario remain below the corresponding QUANTEC 2010 reference limits for bone marrow and whole body. Tumor-to-normal-tissue therapeutic indices are correspondingly favorable under the modelled uptake assumptions.

These dosimetric results are strictly conditional upon the phenomenological uptake fractions assigned to the framework and carry explicit uncertainty of approximately ±50 %. The QUANTEC comparison is implemented solely as a per-cycle safety screen; cumulative multi-cycle claims would require an explicit time-resolved marrow-recovery model that is outside the present scope. Nonetheless, the computational findings indicate that, should the gate condition be satisfied and differential uptake confirmed, the intrinsic radiation characteristics of ❑^28^Mg are compatible with clinically acceptable normal-tissue constraints.

### IV.6. Whole-body and therapeutic-window implications

At the organismal scale the reduction in cellular fitness produced by the Quadruple-Kill cascade is projected onto tumor-burden trajectories generated by the intrinsic Gompertz formulation. The interval between early detectability and the critical transition threshold *N* / (*αK*) = 1 / *e* defines an Intrinsic Therapeutic Window whose temporal width is modulated by the magnitude of *λ_eff_* and by the efficiency of intracellular radio-cofactor delivery.

Because the same magnesium-transport machinery that elevates *θ*_28_ inside malignant cells also operates, albeit at lower intensity, in normal tissues, whole-body implications remain an essential consideration. The framework therefore treats differential uptake as a phenomenological parameter rather than an absolute selectivity claim. Future refinement of this parameter through quantitative biodistribution studies will determine whether the predicted therapeutic window can be exploited without compromising systemic magnesium homeostasis or normal-tissue integrity.

### IV.7. Advantages, limitations and testability of the framework

The principal conceptual advantage of the present approach is the elevation of the Gate Condition from a qualitative premise to a quantitatively linked, continuous variable that can be related to measurable tumor-growth kinetics and transporter activity. By embedding nuclear-decay physics inside endogenous enzymatic architecture, the framework generates emergent multi-scale behaviors (Atomic Switch / DIOC, EDI, Quadruple-Kill, Quintax) that are not immediately apparent from any single organizational level.

Several limitations must be stated explicitly. First, successful catalytic-site occupancy remains an untested input assumption; the chemical-identity approximation 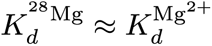 has not been verified experimentally for this isotope. Second, the effective transport coefficient *Φ_transport_* is phenomenological and awaits calibration against measured uptake kinetics. Third, all higher-scale predictions inherit the uncertainty of the biological parameters that lack direct experimental measurement for ^28^Mg. Fourth, the framework is intentionally exploratory rather than predictive of clinical efficacy.

These limitations simultaneously define a clear experimental roadmap. Priority studies include (i) cell-free enzyme assays that correlate EDI metrics with measured catalytic activity, (ii) quantification of differential magnesium uptake and intracellular distribution using isotopically enriched non-radioactive (^25^Mg ICP-MS and ^25^Mg NMR), (iii) single-dose xenograft biodistribution of ^28^Mg, and (iv) multi-cycle toxicity assessment with hematologic monitoring. Only after these decision points are resolved can the computational hypotheses be advanced to more complex in-vivo efficacy models.

### IV.8. Challenges and future experimental priorities

The most immediate challenge is experimental verification of the gate condition itself. Because ^28^Mg is chemically identical to stable magnesium prior to decay, conventional ligand-binding assays cannot distinguish the two species; specialized approaches that combine isotopic enrichment, high-resolution mass spectrometry and real-time catalytic readout will be required.

A second practical challenge is the short physical half-life of ^28^Mg (20.915 h). This constrains both production logistics and the temporal window available for biodistribution, imaging and dosimetry studies. Reliable supply therefore requires either proximity to an existing nuclear-reactor or cyclotron facility capable of producing ^28^Mg (via ^27^Al(α,3p)^28^Mg or ^26^Mg(α,2p)^28^Mg reactions) or the installation of a dedicated compact accelerator within or adjacent to the clinical centre. Under these conditions the isotope can be delivered and administered within 1–2 half-lives of end-of-bombardment, preserving sufficient activity for both imaging and therapy while minimising decay losses during transport and quality-control procedures.

Future work should therefore proceed in carefully staged phases: first establishing preferential uptake and intracellular localisation with stable isotopes, then progressing to single-dose and multi-dose radioactive studies only after the upstream molecular and cellular decision points have been cleared. Parallel refinement of the multiscale computational platform—particularly the incorporation of measured transporter kinetics and time-resolved marrow-recovery dynamics—will allow the framework to evolve from a hypothesis-generating scaffold into a quantitatively predictive tool.

### IV.9. Concluding remarks

The multiscale computational framework presented here explores the logical consequences of treating an essential enzymatic cofactor as an endogenous carrier of radionuclide activity. By making the Gate Condition explicit and by embedding nuclear-decay physics inside endogenous magnesium-dependent architecture, the model generates a coherent cascade of emergent behaviors that span six organizational levels. These behaviors remain strictly conditional upon experimental confirmation of catalytic-site occupancy and differential transport. Should that confirmation be obtained, the radio-cofactor concept would constitute a distinct therapeutic modality—one that acts at the catalytic core of cellular metabolism rather than at downstream signaling or extracellular recognition structures. The present work is offered as a quantitative roadmap intended to reduce mechanistic uncertainty and to accelerate the transition from theoretical proposal to rigorous empirical interrogation.

## V. Conclusion

The present work develops a multiscale computational framework to examine the logical consequences of the Radio-Cofactor Hypothesis, in which an essential enzymatic cofactor itself serves as an endogenous carrier of radionuclide activity. By treating non-zero catalytic-site occupancy by ^28^Mg as an explicit gate condition and by propagating nuclear-decay physics through molecular, organellar, cellular, tissue and whole-body scales, the framework generates a coherent set of emergent behaviors—most notably the Atomic Switch / Decay-Induced Octahedral Collapse, the Enzyme Disruption Index, the Quadruple-Kill cascade and the Quintax Functional Model.

These constructs remain strictly conditional upon experimental verification of the gate condition and of differential magnesium transport. They do not constitute claims of clinical efficacy. Their principal value lies in converting a qualitative therapeutic concept into a quantitatively articulated, experimentally addressable set of mechanistic hypotheses. Should the upstream molecular and transport assumptions be confirmed, the radio-cofactor approach would represent a distinct modality that intervenes at the catalytic core of cellular metabolism rather than at downstream signaling pathways or extracellular recognition structures. The computational roadmap provided here is intended to reduce mechanistic uncertainty and to guide the systematic empirical evaluation required to advance the concept from theoretical proposal to rigorous experimental testing.

## Acknowledgements

The author sincerely thanks IT Engineer Trieu Cao Nguyen for developing the computational engine, executing the simulation framework, and resolving software issues identified during iterative model refinement, thereby enabling the generation of the computational results presented in this study. The author also gratefully acknowledges Dr. Vu Thiên Y for carefully reading the manuscript and providing valuable comments and constructive suggestions that improved its clarity and scientific presentation.

## Declaration of Generative AI and AI-Assisted Technologies in the Manuscript Preparation Process

During the preparation of this manuscript, the author used AI-assisted tools, including *ChatGPT (OpenAI), Gemini (Google), Grok (xAI), and Claude (Anthropic)*, for language refinement, structural editing, and improvement of manuscript readability. All scientific concepts, hypotheses, computational models, interpretations, analyses, and conclusions are solely those of the author. After using these AI-assisted tools, the author critically reviewed, revised, and approved the final manuscript and accepts full responsibility for its content.

## Ethics Statement

This theoretical and computational study involved no human participants, animals, human tissue, identifiable personal data, or clinical interventions. Therefore, ethical approval and informed consent were not required.

