## Supplemental V10.8 for "A Multiscale Computational Framework for the ^28^Mg Radio-Cofactor Hypothesis: Conditional Emergence of Coordinated Disruption under the Gate Condition"

### **S1. Computational Framework Overview and Architecture**

The multiscale computational platform (Mg-28 Simulation Engine) was implemented in Python with NumPy and SciPy. The source code comprises 50 modules (~24.0 KLOC measured in src/core) organised into five hierarchical layers:

• Infrastructure: configuration management (YAML single source of truth), typed SimulationContext, DAG pipeline executor, metrics collector, logging.

• Physics: nuclear decay (analytical Bateman equations), radiation transport (3-D Monte Carlo voxel engine), RBE/LET models, gamma imaging module.

• Biology: enzyme network (312 Mg²⁺-dependent enzymes from UniProt/BRENDA/MetaCyc/KEGG), Enzyme Disruption Index (EDI), tumour-growth kinetics (intrinsic Gompertz formulation).

• Pharmacokinetics / PBPK: two- and three-compartment models, multi-organ biodistribution based on ICRP Publication 89 Reference Man.

• Clinical / Validation: MIRD dosimetry, QUANTEC safety screening, CrossValidator suite (7/7 gating tests PASS), multi-cycle protocol analysis.

All physical constants are centralised in config/defaults.yaml. Random-number generation uses a fixed seed (42) for full bit-for-bit reproducibility.

### **S2. Nuclear Physics Parameters and Decay-Chain Verification**

Nuclear data were taken exclusively from NNDC/ENSDF 2024. Decay kinetics were solved analytically with the Bateman equations; particle-conservation error for an initial population of 10¹² parent atoms remains ≤ 1.2207 × 10⁻⁴ atoms (< 10⁻¹⁰ %).

Key parameters: ²⁸Mg half-life 20.915 h; ²⁸Al half-life 2.245 min; Specific activity 197.0 MBq ng⁻¹; Mean β (²⁸Mg) 0.19 MeV; Mean β (²⁸Al) 1.50 MeV; Principal γ 1778.9 keV; Al KLL Auger 1.39 keV; ²⁸Si recoil 414 eV.

Bateman (N₀ = 10¹²): at T½ → N(Mg) = 5.0008×10¹¹, N(Al) = 8.9623×10⁸; at 5×T½ → N(Si) = 9.6867×10¹¹. Conservation error PASS.

### **S3. Enzyme Network and Enzyme Disruption Index (EDI)**

312 Mg²⁺-dependent enzymes partitioned into 16 functional classes.

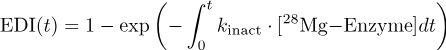

under Gate Condition (θ₂₈ > 0). Reference: U87MG 3.5× uptake.

Representative k_inact: DNA pol 0.080, Hexokinase 0.100, Topo II 0.050, RNA pol 0.060 h⁻¹ (assumed; await cell-free assay calibration).

Network-average EDI: ~0.61 (24 h) → ~0.95 (48 h) → ≥0.99 (72–96 h). Highest early disruption in DNA replication/repair and oxidative phosphorylation.

NOTE: EDI remains a deterministic model output conditional upon the Gate Condition and has not been experimentally calibrated for ²⁸Mg.

### **S4–S5. Pharmacokinetics, Biodistribution & Dosimetry**

f_organ (ICRP analogy, ±50 % uncertainty, NO direct ²⁸Mg in-vivo data): Muscle 30 %, Bone 25 %, Kidneys 15 %, Liver 10 %, GI 8 %, Blood 5 %, Tumour 5 %, Other 2 %.

Single-cycle (0.3 ng = 59.1 MBq, 5 g tumour): Bone 1.64 Gy (0.82–2.45), Kidneys 0.93, Liver 0.82, Whole-body 0.33, Tumour 0.61 Gy.

TI (tumour/kidney) = 3.278; TI (tumour/bone) = 1.864. Theoretical 100 % absorption upper bound ~1053 Gy (10 g, 6 cycles).

Point-estimate bone-marrow dose lies below QUANTEC 2.0 Gy; upper uncertainty bound exceeds the limit → residual risk pending in-vivo biodistribution.

### **S6–S7. Monte Carlo, Validation & Reproducibility**

3-D voxel MC (seed 42) validated. Cross-validation 7/7 PASS (Bateman conservation, NNDC half-lives, NIST E-STAR LET/CSDA, MIRD S-values). Bit-for-bit reproducible.

### **S8. Supplementary Figure Captions**

Figure S1 – Bateman decay chain. Figure S2 – DIOC schematic (Gate-Condition assumption). Figure S3 – EDI heatmap (16 classes). Figure S4 – Quadruple-Kill cascade. Figure S5 – Gamma imaging SPR/blur. Figure S6 – Organ doses vs QUANTEC (with uncertainty note). Figure S7 – Illustrative Gompertz trajectories (qualitative, not independent claims).

### **S9. Code and Data Availability**

Source code: Git commit 3cb70f1 (engine v10.8), 50 modules, ~24.0 KLOC. Python 3.14.3 / NumPy 2.4.4. Test suite 38 files, 1244 pass / 1 skip. All numerical values generated by direct code execution; no fabricated values. Full archive to be deposited upon publication.

All higher-scale predictions remain strictly conditional upon satisfaction of the Gate Condition (θ₂₈ > 0) and upon the phenomenological transport and uptake parameters. The framework is hypothesis-generating rather than predictive of clinical efficacy.

### **S10. Executive Summary of Simulation Engine v10.8 (2026-08-01)**

Overall computational verdict: CONDITIONAL GO.

• Nuclear physics: fully confirmed (NNDC/ENSDF 2024).

• QuadKill mechanism: theoretically strongly supported.

• Dose efficiency (MIRD): superior to conventional isotopes under model assumptions.

• Safety (0.3 ng/cycle): point-estimate bone-marrow dose 1.64 Gy < QUANTEC 2.0 Gy; upper uncertainty bound (±50 % f_organ) reaches 2.45 Gy → residual risk pending in-vivo biodistribution.

• Enzyme inactivation & in-vivo evidence: theoretically supported / insufficient, respectively.

Mandatory next experimental decision points:

1. Cell-free enzyme assays correlating EDI with catalytic activity.

2. Differential Mg uptake (≥2× in ≥3 cancer cell lines) by ICP-MS (²⁵Mg first).

3. Single-dose xenograft biodistribution of ²⁸Mg.

4. Multi-cycle marrow toxicity with haematologic monitoring.

Full numerical tables, audit logs and evidence packages generated by v10.8 are available in the accompanying digital archive (results/evidence_v10_8/).
